# PTMOverlay: A Proteomic Tool to Visualize Post-Translational Modifications Across Evolution

**DOI:** 10.64898/2026.02.03.703592

**Authors:** Camille Krieger, Zachary Everton, Youngki You, Brandon Lewis, Thomas Bank, Meagan C. Burnet, Sarai Williams, Hanna E. Walukiewicz, Christopher V. Rao, Alan J. Wolfe, Samuel H. Payne, Ernesto S. Nakayasu

**Affiliations:** Department of Biology, Brigham Young University, Provo, Utah, USA; Biological Sciences Division, Pacific Northwest National Laboratory, Richland, Washington, USA; Department of Microbiology and Immunology, Stritch School of Medicine, Health Sciences Division, Loyola University Chicago, Maywood, Illinois, USA; Environmental and Molecular Sciences Division, Pacific Northwest National Laboratory, Richland, Washington, USA; Department of Chemical and Biomolecular Engineering, University of Illinois at Urbana-Champaign, Urbana, Illinois, USA

## Abstract

Evolutionary conservation has been considered a hallmark of essential basic functions in cells. Therefore, the study of evolutionarily conserved post-translational modifications (PTMs) can provide insight into their role in protein function. In this context, mass spectrometry can identify and quantify thousands of PTM sites. However, a major bottleneck lies in analyzing the large amounts of data collected by the mass spectrometer. Here we address the need for a protein sequence alignment tool for multiple PTMs across several species. We developed a tool named PTMOverlay that takes peptide identification output files and overlays PTM sites onto multiple protein sequence alignments. Examining 31 bacteria isolates, we combined their protein sequences with select PTM types, including acetylation, phosphorylation, monomethylation, dimethylation, and trimethylation. The tool revealed a variety of conserved modification sites on the bacterial central carbon metabolism. Further structural analysis revealed possible interactions between methylated arginine and lysine residues with phosphothreonine/serine sites on the homodimer interface of enolase. Overall, this tool can parse large amounts of mass spectrometry data and allows for more informed and efficient selection of sites for future studies of protein function.

## Main Text

Post-translational modifications (PTMs) alter the physicochemical properties of amino acid residues, thereby changing protein structure, regulating enzyme activity, stabilizing proteins, and influencing protein interactions and function [1, 2]. Common PTMs include phosphorylation, methylation, and ubiquitination, but over 680 different PTM types have been described [3]. Therefore, the study of PTMs has great potential to enhance the understanding of protein regulation and function. Mass spectrometry has been instrumental for the study of PTMs, often identifying and quantifying thousands of PTM sites in a single experiment, but the analysis of the large dataset is a bottleneck [4].

Evolutionary conservation has been considered a hallmark of selective pressure and can indicate essential function [5]. This same principle can be applied to PTMs, being evolutionarily conserved modifications likely to have an essential function. We have previously demonstrated that conserved acetylation sites on bacterial proteins regulate the central carbon metabolism and the ribosome, while conserved methylation sites on the ribosomal protein bL11 regulate the stringent response to nutrient starvation [6-8]. However, parsing these evolutionarily conserved PTM sites from large mass spectrometry datasets through protein alignment remains a bottleneck.

Here, we developed a computational tool named PTMOverlay for aligning evolutionarily conserved PTMs across multiple species. First, sequences from protein databases of individual species are mapped based on KEGG (Kyoto Encyclopedia of Genes and Genomes) annotations [9] with GhostKOALA [10]. Then, orthologous proteins are submitted to multiple sequence alignment using MUSCLE [11]. Finally, PTMOverlay obtains tandem mass spectra search result pepXML (common format) files and overlays the identified PTMs (**Fig. 1**). To test PTMOverlay, we extracted proteins from a set of 31 bacterial isolates from 29 species (**Tab. 1**), digested them and analyzed them by liquid chromatography-tandem mass spectrometry. The generated datasets were searched for PTMs involving acetylation, carbamylation, phosphorylation, monomethylation, dimethylation, and trimethylation. When queried at a KEGG pathway level, the PTMOverlay tool can provide information on conserved modification sites across the proteins from the same pathway. When applied to the glycolysis/gluconeogensis pathway, PTMs were found in conserved lysine (acetylation, carbamylation and mono/di/trimethylation), arginine (mono and dimethylation) and threonine (phosphorylation) residues from glyceraldehyde 3-phosphate dehydrogenase (GAPDH), phosphoglycerate kinase (PGK), enolase (ENO), phosphofructokinase (PFK), aldehyde-alcohol dehydrogenase (adhE), glucose-6-phosphate isomerase (GPI), pyruvate kinase (PK), fructose-biphosphate aldolase (FBA), triosephosphase isomarase (TPI), and glucokinase (glk) (**Fig. 2**). For all the listed enzymes, PTMs were observed in the conserved amino acid residues from >50% of the species. For instance, K130 of GAPDH was either carbamylated or acetylated in 24 out of the 29 species, while R12 of enolase was mono/di/trimethylated in 23 out of the 29 species.

**Figure 1.**
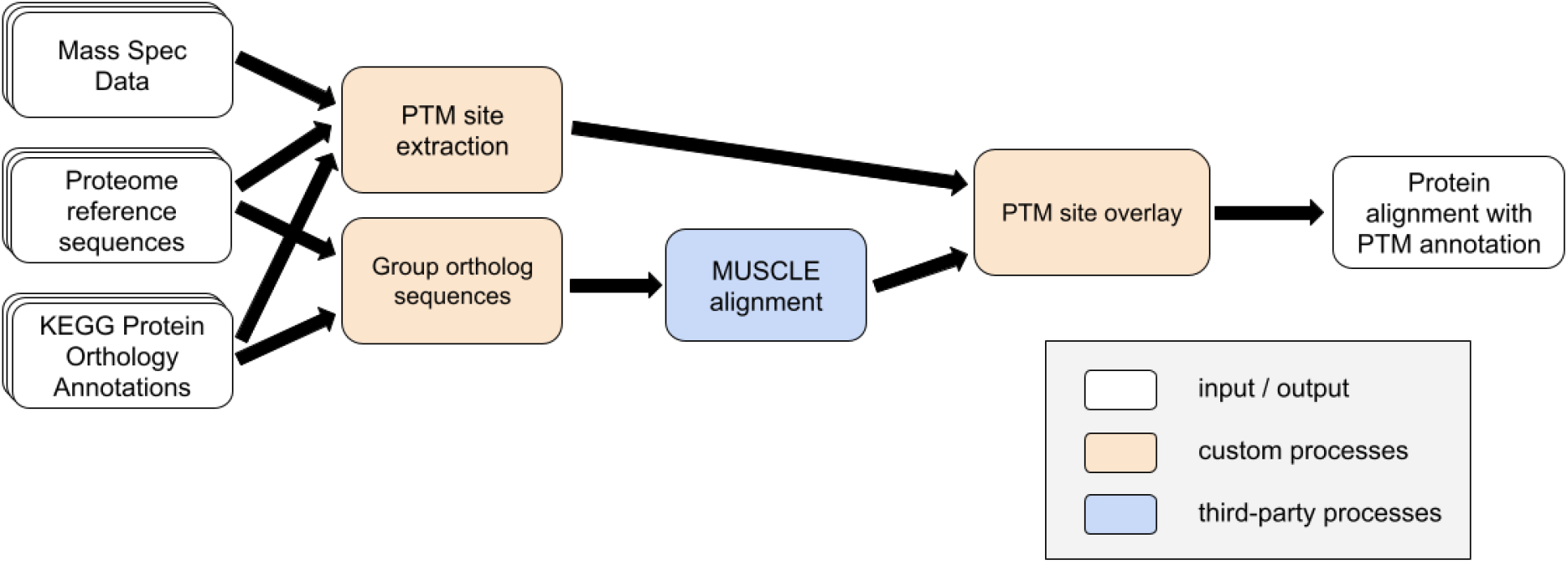
Workflow of PTMOverlay. PTMOverlay requires 3 input files mass spectrometry database searching results in the common pepXML format, individual species/strains sequence database in FASTA format, and KEGG annotations of the sequence databases. This later piece can be annotated using GhostKOALA. PTMOverlay group ortholog sequences based on the KEGG annotation, which are then aligned using MUSCLE. Then the PTM sites are extracted from pepXML and overlayed onto the multiple sequence alignment.

**Table 1.**
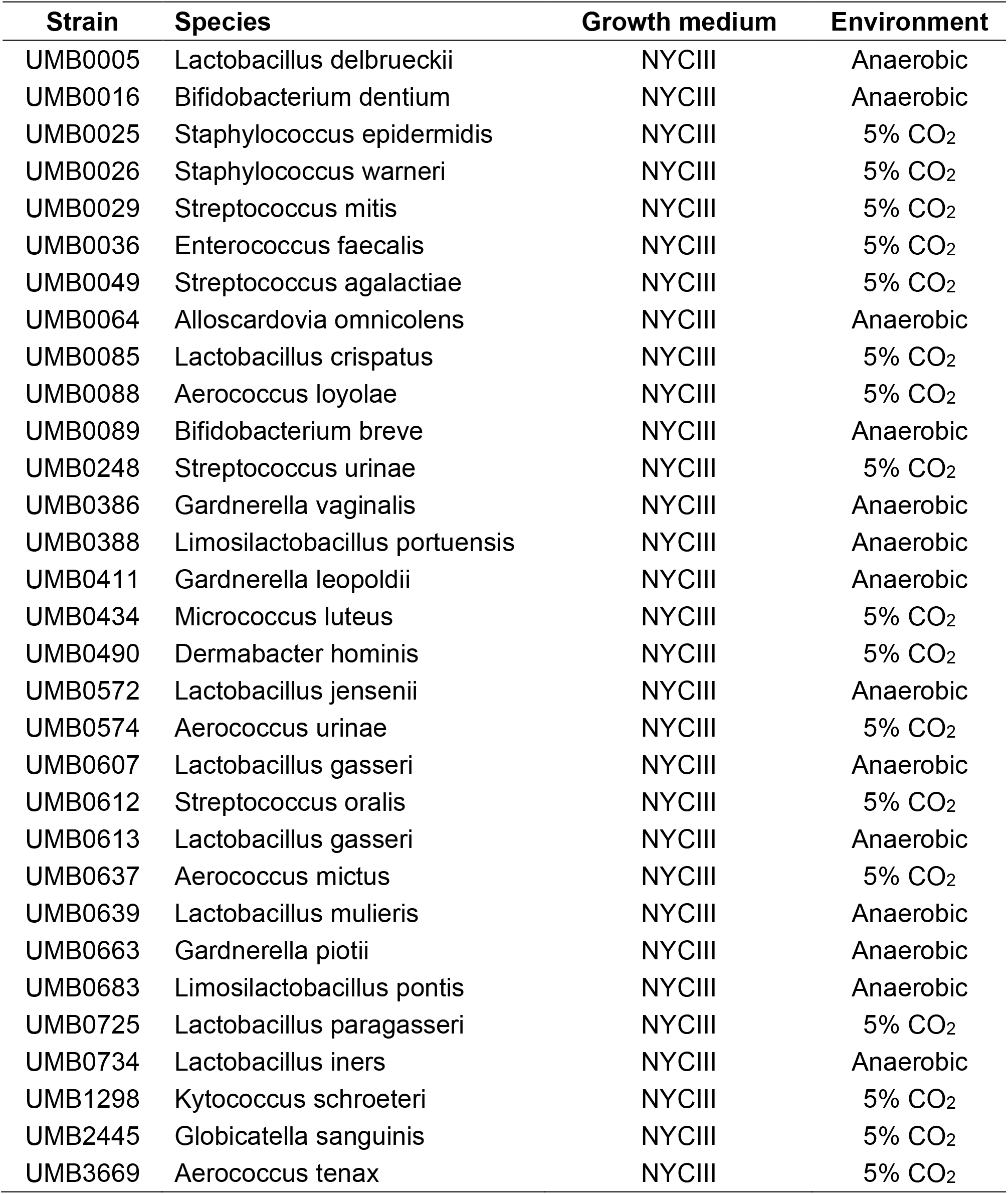
Urinary bacterial isolates used in this study with incubation conditions.

**Figure 2.**
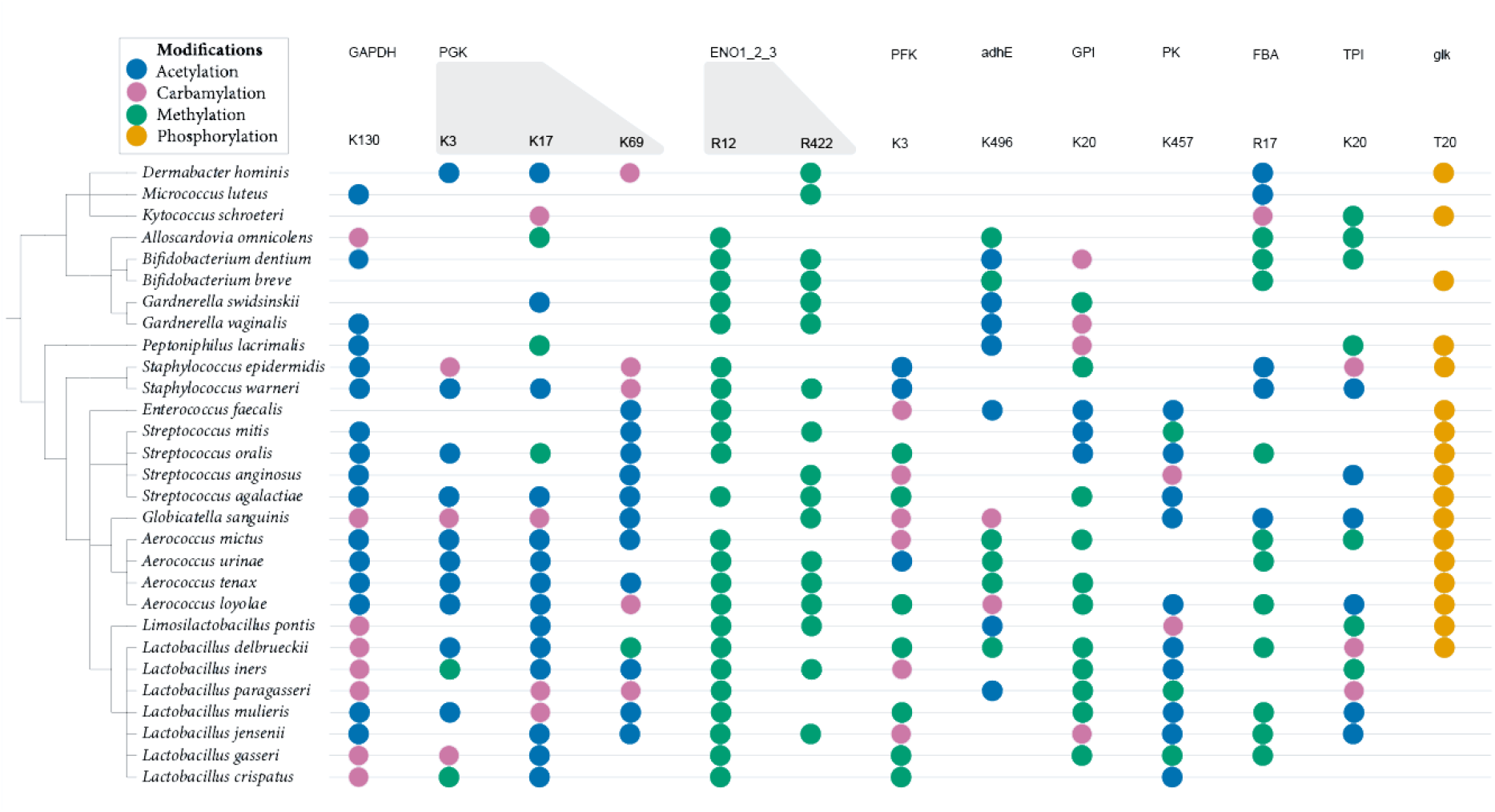
Conserved post-translational modifications within the glycolysis/gluconeogenesis pathway. The dot plot graph represents invariant amino acid residues on proteins from the glycolysis/gluconeogenesis pathway that are modified by acetylation, carbamylation, methylation (combining mono, di and tri), and phosphorylation. The amino acid residue number is based on the *Dermabacter hominis* sequence.

PTMOverlay also provides easy visualization of conserved sequences along with the modification sites. For example, on the N-terminal region of enolase, at position 5 (based on *Streptococcus mitis* sequence) there is a partially conserved threonine phosphorylation site (**Fig. 3**). The remaining sequences had acidic amino acids, glutamate or aspartate, at the same or adjacent positions (**Fig. 3**), suggesting that a negative charge (from the acidic amino acid side chain or phosphate group) might play an important function in this protein. Downstream at position 8, 34 out of 35 sequences (some species have multiple isoforms) have an aromatic amino acid, either histidine or tyrosine, with the tyrosine residue phosphorylated in 9 of the species (**Fig. 3**). At position 10, an arginine residue is conserved in 31 out of the 35 sequences and is mono or dimethylated in 24 of the sequences (**Fig. 3**). At position 190, an invariant lysine residue is acetylated in 8 sequences, while 1 sequence is dimethylated and 1 sequence is carbamylated. On the C-terminal region, at arginine 403 is 100% conserved and mono or dimethylated in 18 of the sequences, suggesting another possible conserved function.

**Figure 3.**
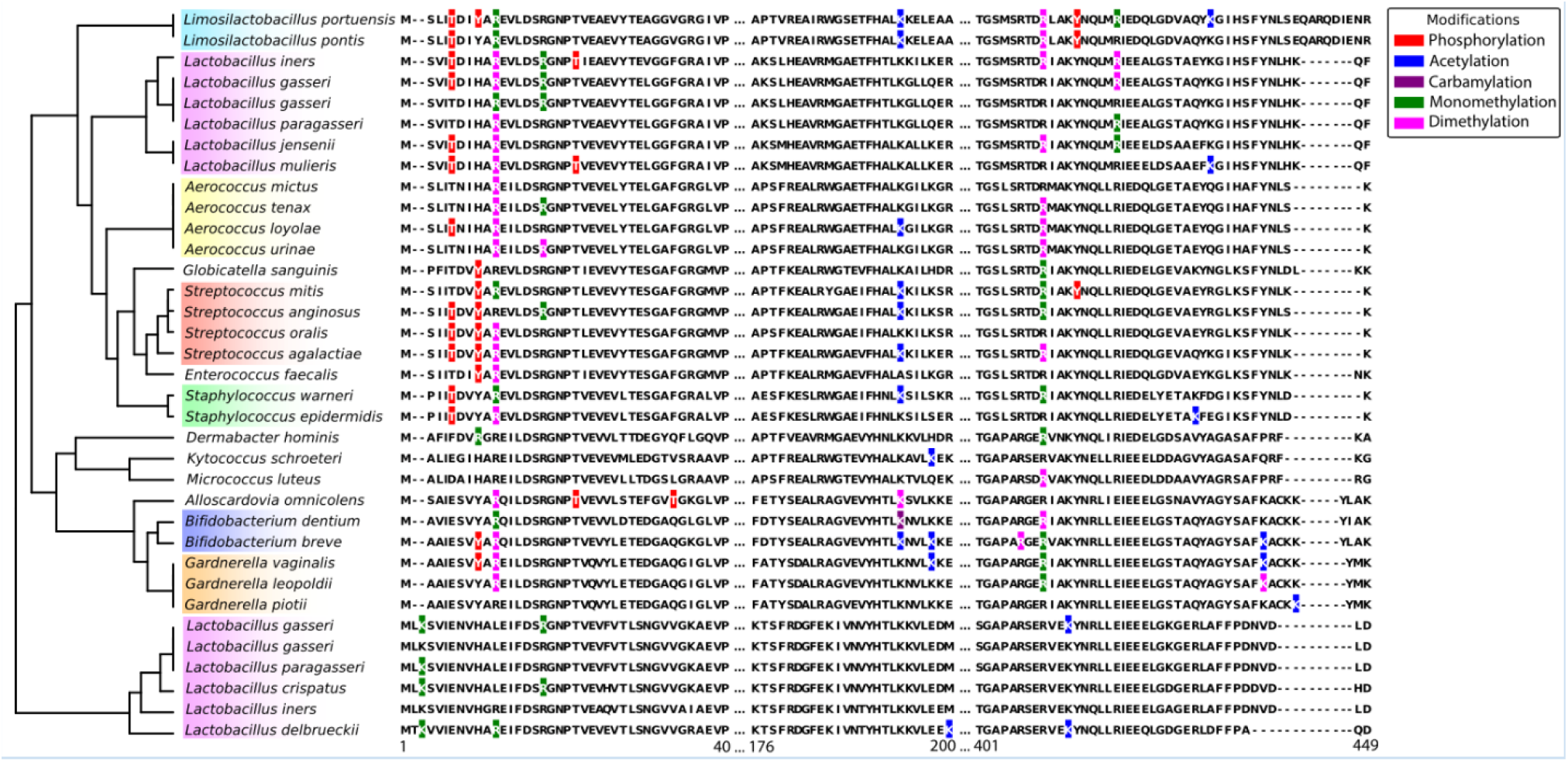
Post-translational modification conservation in enolase across bacteria. The multi-sequence alignment shows enolase regions with conserved PTMs. The residue numbers are from the PTMOverlay result after multi-sequence alignment. Some strains have two enolase isomers. Phosphorylation (red), acetylation (blue), carbamylation (purple), monomethylation (green), dimethylation (magenta)

To further select specific sites to study, we often analyze the structure of the protein, searching for modifications that would impact the structure or activity of the protein. As an example, we analyzed the 3D structure of the enolase from *Streptococcus pneumoniae (*Uniprot Q97QS2), which has identical sequence to *S. mitis. S. pneumoniae* enolase is composed of 4 dimers that are assembled into an octamer (**Fig. 4**), similar to enolases of other related species [12-14]. All the conserved modifications mentioned above occur on amino acid residues within or close to the dimer interface (**Fig. 4**). Threonine 5 is on the N-terminal tail (**Fig. 4**) and its phosphorylation is unlike to interfere with the dimer stability. The N-terminal tail is parallel to the active site. Therefore, it is possible that threonine 5 phosphorylation has a role in regulating the enzymatic activity. In *E. coli*, enolase phosphorylation stabilizes its otherwise reversible activity to convert 2-phosphoglycerate into phosphoenolpyruvate [15]. Unfortunately, the phosphorylation site responsible for this regulation has not been mapped to enable us to compare if *S. mitis* threonine 5 would have similar activity. The acetylation of lysine 190 might impact dimer stability. The positive charge of lysine 190 and the negative charge of aspartate 204 are positioned close enough to potentially form a salt bridge, with 4.4 Å, which may reduce to 3.2 Å due to potential side chain dynamics. However, after acetylation, lysine loses its positive charge, and the acetyl group would be at only 2.5 Å from aspartate 204 (**Fig. 4A**). The modification would break the salt bridge and may cause a steric conflict between the two residues. The mono- or di-methylation on arginine 403 might also have an impact on the enolase dimer stability (**Fig. 4B**). Arginine 403 forms five hydrogen bonds with aspartate 204, arginine 372, aspartate 402, and tyrosine 407. It also forms a salt bridge with the negatively charged aspartate 402. The amphipathic chain of arginine 403 facilitates hydrophobic interactions with arginine 400, which has multiple interactions with residues from the other monomer, thereby facilitating enolase dimerization [12]. Mono- or dimethylation could alter the various interactions between arginine 403 and the other residues, while the enlarged side chain of arginine 403 could potentially lead to steric conflicts with nearby residues. Lastly, phosphorylation on tyrosine 8 and monomethylation on arginine 10 might similarly influence enolase dimer stability. Tyrosine 8 and arginine 10 are close together with aspartate 415 from the adjacent chain (**Fig. 4C**). Arginine 10 monomethylation would make the residue more hydrophobic and increase the interaction with tyrosine 8 (**Fig. 4C**), possibly helping to stabilize the dimer. Conversely, tyrosine 8 phosphorylation would cause a structure change due to the lack of space (**Fig. 4C**). Furthermore, the negative charge of the phosphate would repulse the negative charge of the aspartate 415 further changing the structure and possibly causing disruption of the dimer. Therefore, in a scenario that the goal is to study the dimer stability tyrosine 8 phosphorylation and arginine 10 monomethylation would be excellent targets to study, while would be a candidate for studying enzymatic activity regulation.

**Figure 4.**
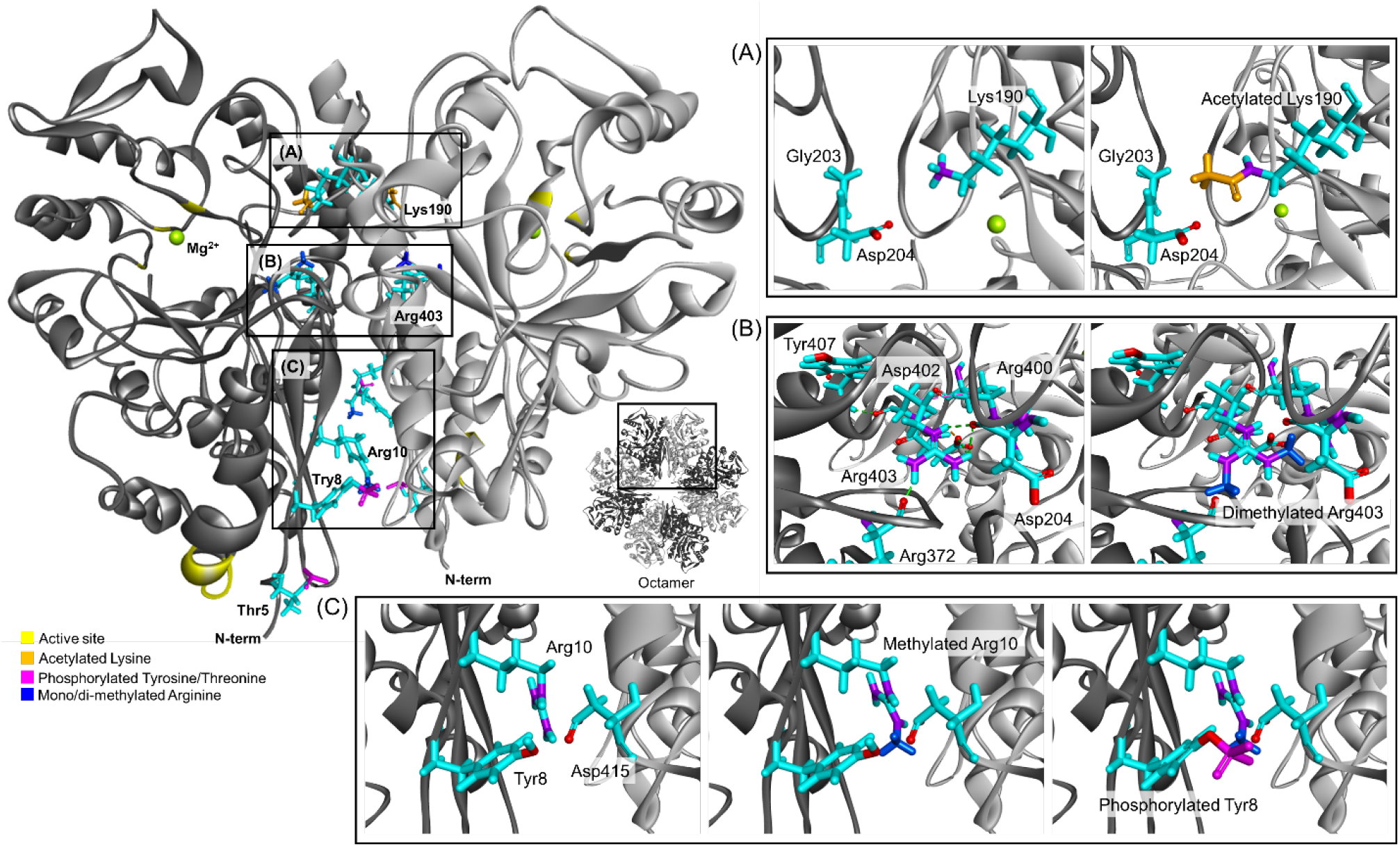
Conserved post-translational modifications on the dimer interface of enolase (*Streptococcus mitis*). (A) Lys-190 is 4.4 Å closer to Asp-204 after acetylation of Lys-190. (B) Arg-403 forms five hydrogen bonds with Asp-204, Arg-372, Asp-402, and Tyr-407 (green colored line). There is a salt bridge between Arg-403 and Asp-402 (orange colored line). Arg-403 forms hydrophobic interactions with Arg-400 (magenta colored line). (C) Tyr-8, Arg-10, and Asp-415 are located within 4.2 Å without post-translational modification. Oxygen and nitrogen atoms are colored red and violet, respectively. The structure displayed is the crystal structure of enolase from *Streptococcus pneumoniae* (PDB 1W6T).

These results demonstrate the power of the PTMOverlay tool for helping to interpret possible biological roles of PTMs from large-scale proteomic datasets. While enolase and glycolysis served as examples, the tool is fully customizable and can be applied to any set of proteins or pathways with KEGG annotations. Users can specify protein sets or pathways and adjust the level of sequence conservation. This flexibility enables researchers to efficiently screen broad datasets, identify modification sites of interest, and prioritize targets for downstream biochemical or functional studies. Although numerous algorithms exist for protein sequence alignment [13-15], relatively few incorporate PTM information directly into the aligned residues, and those that do typically focus on only a single PTM type [16]. Overlaying multiple PTMs opens the opportunity to study PTM crosstalk [17]. Therefore, alignments that integrate multiple PTM types offer more informative comparisons across species and may provide additional insights into understanding the complexity of protein regulation.

Another aspect we considered when developing PTMOverlay was the file formats. Database searching tools for peptide and PTM identification use a wide range of file formats, leading to problems in software compatibility [18]. Considering this potential issue, we developed PTMOVerlay to accept the common pepXML format [19]. Not all database searching tools have the pepXML format as an option for peptide identification result output. However, results from many different database searching tools can be easily converted to pepXML with IDFileConverter using OpenMS or ProteoWizard [20, 21]. Therefore, the use of the pepXML common for should facilitate the adoption of PTMOverlay by users.

In summary, here we developed a PTM site alignment tool that uses the common pepXML peptide identification files and parse multiple types of PTMs. PTMOverlay enables the study of PTMs not only in the context of evolutionary conservation but also in the context of possible crosstalk with adjacent or competing modifications. We further provide an example on how the combination of PTMOverlay with protein structural analysis can give additional insights and help prioritize sites for further investigation.

## Experimental Procedures

### Strains

Urinary isolates for testing were procured from University of Loyola Chicago IRB-approved biorepository (LU 215192). A complete list of strains used in this study can be found in **Table 1**. Each strain tested was grown from frozen stock, frozen in Brucella media at −80°C. Strains were struck out on trypticase soy agar (TSA) with 5% sheep blood, and identity was confirmed via MALDI-TOF MS.

### Bacterial Culture Conditions and Biomass Harvesting

NYCIII broth was prepared as outlined for ATCC medium no. 1685 with the following modification: 4.2 mL HEPES buffer solution, 7.5 g proteose peptone no. 3, 2.5 g glucose, 2.5 g NaCl, and 1.875 g yeast extract were added to 450 mL Millipore water and autoclaved at 121 °C for 15 min. After cooling, 25 mL heat-inactivated horse serum and 25 mL heat-inactivated newborn calf serum were added by vacuum filtration (0.2 μm) to yield a final volume of 500 mL. Single colonies of test strains were inoculated into 5 mL of this NYCIII broth in culture tubes and incubated for 24 h at 37 °C either anaerobically or in ambient atmosphere supplemented with 5% CO_2_, with shaking at 200 rpm (Tab 1). Cultures were then transferred into 25 mL NYCIII broth in Erlenmeyer flasks for biomass expansion and incubated for an additional 48 h under the same conditions. Cells were harvested by centrifugation at 2,000 rpm for 30 min, washed twice with deionized water, and stored at −80 °C.

### Protein digestion

Cell pellets were resuspended in 100 mM NH_4_HCO_3_ and transferred to 2-mL tubes containing 0.5 mm glass beads (OMNI International, 2 mL Bead Kit, 0.5 mm glass beads; cat. 19-622). Cells were lysed by vortexing 6 times for 30 s, with 30 s incubations on ice between vortex cycles. The lysates were cooled on ice, and a 25-gauge needle was used to puncture the bottom of each 2 mL tube, which was then placed into a 4-mL cryovial and centrifuged at 1,000 × g for 10 min at 4 °C to separate the lysate from the glass beads. Protein concentration was determined by BCA assay (Thermo Fisher Scientific), and an aliquot of 500 µg protein was submitted to digestion. One volume of 5% w/v sodium deoxycholate, 100 mM dithiothreitol in 100 mM NH_4_HCO_3_ was added to nine volumes of samples, which were denatured at 70 °C for 30 min in a thermomixer at 850 rpm, then rapidly cooled on ice. Sequencing-grade trypsin was then added at a 1:50 (w/w) enzyme:protein ratio, and samples were incubated for 3 h at 37 °C at 850 rpm. Digestion was stopped by addition of trifluoroacetic acid to a final concentration of 1% (v/v). Samples were centrifuged at 10,000 × g for 5 min at room temperature to remove debris and desalted using C18 solid-phase extraction cartridges (Phenomenex, 50 mg/1 mL) according to the manufacturer’s instructions. Eluates were snap-frozen in liquid nitrogen and concentrated down to 50 µL in a vacuum centrifuge. Peptide concentration was measured by BCA assay and samples were diluted to 0.1 µg/µL in milliQ water.

### Mass spectrometry and data analysis

Peptides were dissolved in 2% acetonitrile and 0.1% trifluoroacetic acid and separated on an in-house–packed reversed-phase analytical column (25 cm length, 360 μm o.d. × 75 μm i.d. fused silica picofrit capillary, New Objective) packed with ReproSil-Pur 120 C18-AQ, 1.9 μm stationary phase. The column was connected to a nanoACQUITY UPLC system (Waters) and maintained at 50 °C using an AgileSLEEVE column heater (Analytical Sales and Services). Peptides were eluted using a linear gradient from 8% to 35% buffer B (0.1% formic acid in acetonitrile) over 110 min at a flow rate of 200 nL/min. Electrospray ionization was performed in positive ion mode using a Nanospray (NSI) source with a spray voltage of 2.2 kV and an ion transfer tube temperature of 300 °C. Peptides were analyzed on an Orbitrap Fusion Lumos mass spectrometer (Thermo Fisher Scientific, serial number FSN20129) operated with FAIMS in three compensation voltage settings (−40, −60, and −80 V). For each FAIMS CV, full MS1 scans were acquired in the Orbitrap at a resolution of 120 K over an m/z range of 300–1800 with an AGC target of 4 × 10^5^ and a maximum injection time of 50 ms. Data-dependent MS2 spectra were acquired in the ion trap using quadrupole isolation (1.6 m/z window) and HCD fragmentation with a normalized collision energy of 32%, an AGC target of 1 × 10^4^, and a maximum injection time of 35 ms. Precursors with charge states 2–7 and intensities above 5,000 were selected for fragmentation, with dynamic exclusion of previously sequenced precursors for 60 s within a ±10 ppm window. Data were processed with MSFragger (v4.0) using FragPipe (v21.1), searching against each species’ reference proteome database from GenBank (downloaded on January 27, 2025). Search parameters included protein N-terminal acetylation and methionine oxidation as variable modifications, and carbamidomethylation of cysteine residues as a fixed modification. For phosphoproteome analysis, phosphorylation of serine, threonine, and tyrosine residues was specified as a variable modification. Acetylation and carbamylation of lysine residues were also set as variable modifications with previously reported parameters. For methylation, mono- and di-methylation of lysine and arginine residues, as well as tri-methylation of lysine residues, were included as variable modifications. All PTM types were searched independently.

### Post-Translational Modification Overlay and Analysis

PTMs for each bacterial species were extracted from the raw mass spectrometry files. Protein sequences were obtained from corresponding reference proteome files. Using these sequences, orthologous proteins were gathered searching the [KEGG](https://www.genome.jp/kegg/) database. Proteins were retained for downstream analysis only if sequences were available from at least two species. Multiple sequence alignments (MSAs) were performed using [MUSCLE](https://drive5.com/muscle/downloads_v3.htm) accessed through a web-based application programming interface (API). For each protein, an HTML file was generated displaying the MSA with the PTMs overlaid. Phylogenetic analyses were performed on the aligned sequences using the Bio.Phylo.TreeConstruction package. The resulting trees were exported, stylized in Adobe Illustrator, and appended to the MSA visualization. For consistency, the aligned sequences used in phylogenetic inference were parsed directly from the HTML alignment files, including the annotated PTM positions. To extend the pipeline beyond individual protein analyses, we included features to identify highly conserved PTMs across entire pathways. Using the annotated alignments, KEGG ids, and the Pandas package, CSV files were generated displaying the species and each PTM site along the protein pathway. PTM sites shared by at least 50% of species were classified as highly conserved and included in pathway-level visualizations. Phylogenetic trees for pathway-level comparisons were constructed using the ncbi-taxonomist package (https://github.com/ridgelab/NCBI-Taxonomy-Builder) and visualizations were generated using the svg_stack package (https://github.com/astraw/svg_stack/tree/main). All pipeline parameters are user-configurable through a centralized configuration file. Users may specify the minimum number of species required for inclusion in an MSA, define custom protein sets or pathways of interest, and adjust all analysis thresholds. This ensures the workflow remains flexible, reproducible, and scalable across diverse protein families. The entire workflow is implemented as a Snakemake pipeline with a Docker container, enabling reproducible execution from a single command.

## Code Availability

Project code is available at GitHub at URL: https://github.com/evergreen700/PTMOverlay

## Acknowledgements

Work was performed in the Environmental Molecular Science Laboratory, a U.S. Department of Energy (DOE) national scientific user facility at Pacific Northwest National Laboratory (PNNL) in Richland, WA. Battelle operates PNNL for the DOE under contract DE-AC05-76RLO01830. This work received supported by the U.S. Department of Energy, Office of Science, Office of Biological and Environmental Research, Early Career Research Program (S.H.P.). E.S.N. was partially supported by the National Institute of Diabetes and Digestive and Kidney Diseases (R01 DK138335).

## References

[1] Bilbrough, T., Piemontese, E., Seitz, O., Dissecting the role of protein phosphorylation: a chemical biology toolbox. Chem Soc Rev 2022, 51, 5691–5730.

[2] Muller, M. M., Post-Translational Modifications of Protein Backbones: Unique Functions, Mechanisms, and Challenges. Biochemistry 2018, 57, 177–185.

[3] Huang, X., Feng, Z., Liu, D., Gou, Y., et al., PTMD 2.0: an updated database of disease-associated post-translational modifications. Nucleic Acids Res 2025, 53, D554–D563.

[4] Olsen, J. V., Mann, M., Status of large-scale analysis of post-translational modifications by mass spectrometry. Mol Cell Proteomics 2013, 12, 3444–3452.

[5] Jordan, I. K., Rogozin, I. B., Wolf, Y. I., Koonin, E. V., Essential genes are more evolutionarily conserved than are nonessential genes in bacteria. Genome Res 2002, 12, 962–968.

[6] Feid, S. C., Walukiewicz, H. E., Wang, X., Nakayasu, E. S., et al., Regulation of Translation by Lysine Acetylation in Escherichia coli. mBio 2022, 13, e0122422.

[7] Nakayasu, E. S., Burnet, M. C., Walukiewicz, H. E., Wilkins, C. S., et al., Ancient Regulatory Role of Lysine Acetylation in Central Metabolism. mBio 2017, 8.

[8] Walukiewicz, H. E., Farris, Y., Burnet, M. C., Feid, S. C., et al., Regulation of bacterial stringent response by an evolutionarily conserved ribosomal protein L11 methylation. mBio 2024, 15, e0177324.

[9] Kanehisa, M., Furumichi, M., Tanabe, M., Sato, Y., Morishima, K., KEGG: new perspectives on genomes, pathways, diseases and drugs. Nucleic Acids Res 2017, 45, D353–D361.

[10] Kanehisa, M., Sato, Y., Morishima, K., BlastKOALA and GhostKOALA: KEGG Tools for Functional Characterization of Genome and Metagenome Sequences. J Mol Biol 2016, 428, 726–731.

[11] Edgar, R. C., MUSCLE: multiple sequence alignment with high accuracy and high throughput. Nucleic Acids Res 2004, 32, 1792–1797.

[12] Hosaka, T., Meguro, T., Yamato, I., Shirakihara, Y., Crystal structure of Enterococcus hirae enolase at 2.8 A resolution. J Biochem 2003, 133, 817–823.

[13] Randic, M., Pisanski, T., Protein alignment: Exact versus approximate. An illustration. J Comput Chem 2015, 36, 1069–1074.

[14] Sievers, F., Wilm, A., Dineen, D., Gibson, T. J., et al., Fast, scalable generation of high-quality protein multiple sequence alignments using Clustal Omega. Mol Syst Biol 2011, 7, 539.

[15] Thompson, J. D., Higgins, D. G., Gibson, T. J., CLUSTAL W: improving the sensitivity of progressive multiple sequence alignment through sequence weighting, position-specific gap penalties and weight matrix choice. Nucleic Acids Res 1994, 22, 4673–4680.

[16] Miller, M. L., Blom, N., Kinase-specific prediction of protein phosphorylation sites. Methods Mol Biol 2009, 527, 299-310, x.

[17] Leutert, M., Entwisle, S. W., Villen, J., Decoding Post-Translational Modification Crosstalk With Proteomics. Mol Cell Proteomics 2021, 20, 100129.

[18] Deutsch, E. W., File formats commonly used in mass spectrometry proteomics. Mol Cell Proteomics 2012, 11, 1612–1621.

[19] Keller, A., Eng, J., Zhang, N., Li, X. J., Aebersold, R., A uniform proteomics MS/MS analysis platform utilizing open XML file formats. Mol Syst Biol 2005, 1, 2005 0017.

[20] Chambers, M. C., Maclean, B., Burke, R., Amodei, D., et al., A cross-platform toolkit for mass spectrometry and proteomics. Nat Biotechnol 2012, 30, 918–920.

[21] Rost, H. L., Sachsenberg, T., Aiche, S., Bielow, C., et al., OpenMS: a flexible open-source software platform for mass spectrometry data analysis. Nat Methods 2016, 13, 741–748.

